## Supplementary for "Robust antibiotic sensitization of pathogenic *Pseudomonas aeruginosa* via negative hysteresis in the cell envelope"

### Title:

* Shared first or last author

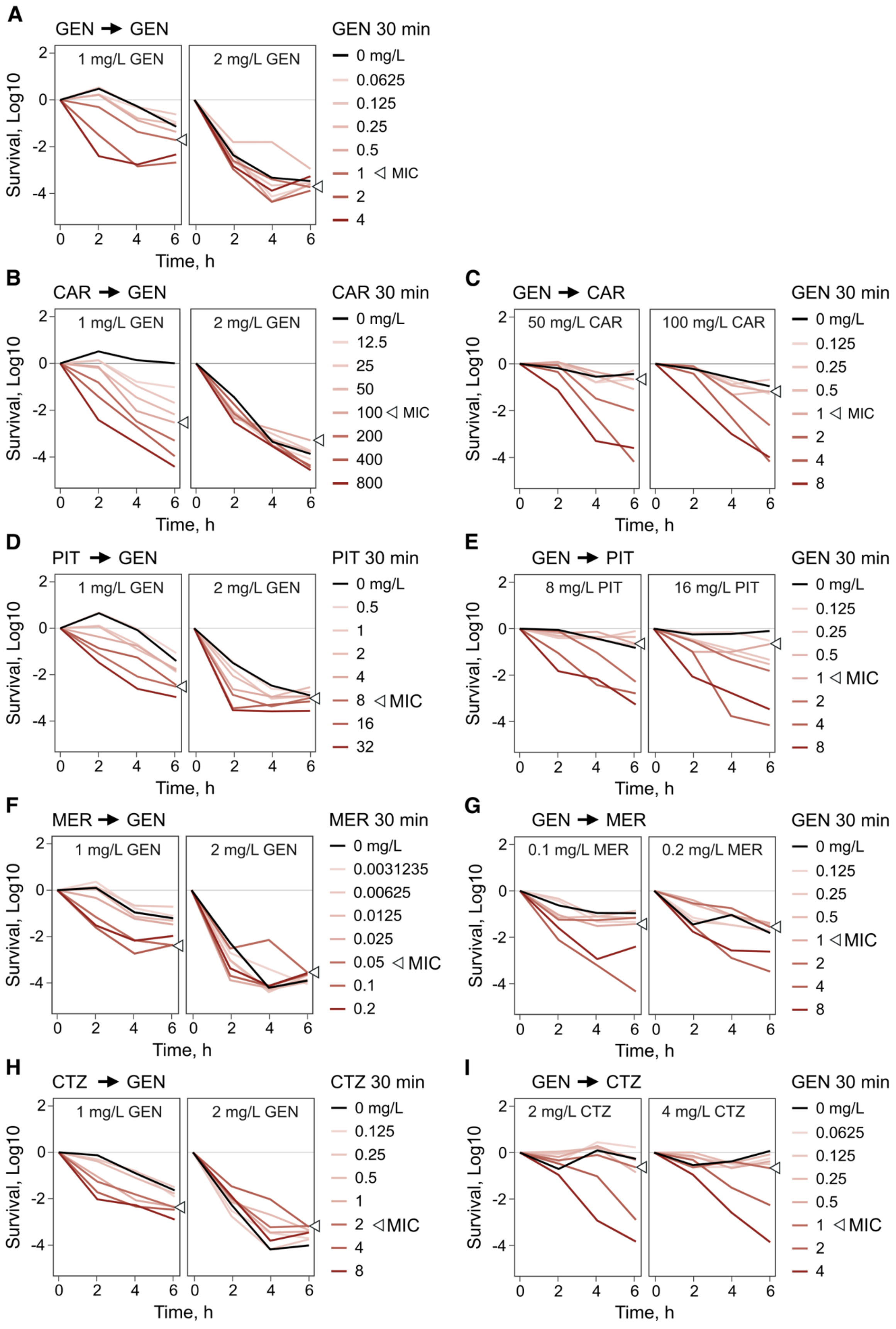
A
GEN → GEN
1 mg/L GEN
2 mg/L GEN
GEN 30 min
0 mg/L
0.0625
0.125
0.25
0.5
1 ◁ MIC
2
4
Survival, Log10
Time, h
B
CAR → GEN
CAR 30 min
0 mg/L
12.5
25
50
100 ◁ MIC
200
400
800
C
GEN → CAR
50 mg/L CAR
100 mg/L CAR
GEN 30 min
0 mg/L
0.125
0.25
0.5
1 ◁ MIC
2
4
8
D
PIT → GEN
PIT 30 min
0 mg/L
0.5
1
2
4
8 ◁ MIC
16
32
E
GEN → PIT
8 mg/L PIT
16 mg/L PIT
GEN 30 min
F
MER → GEN
MER 30 min
0 mg/L
0.0031235
0.00625
0.0125
0.025
0.05 ◁ MIC
0.1
0.2
G
GEN → MER
0.1 mg/L MER
0.2 mg/L MER
GEN 30 min
H
CTZ → GEN
CTZ 30 min
0 mg/L
0.125
0.25
0.5
1
2 ◁ MIC
4
8
I
GEN → CTZ
2 mg/L CTZ
4 mg/L CTZ
GEN 30 min
0 mg/L
0.0625
0.125
0.25
0.5
1 ◁ MIC
2
4

36

**Fig. S1: Detailed dose-dependency of various hysteresis interactions with gentamicin (GEN). A:** Time-kill curves after 30 min GEN pre-treatment followed by GEN main treatment. **B:** Time-kill curves after 30 min CAR pre-treatment at various concentrations followed by GEN main treatment at two concentrations. The hysteresis effect increased with increasing concentration of the pre-treatment antibiotic and to a larger extent than observed for GEN pretreatments (left plot). At a higher main treatment concentration (right plot), high killing rates were independent from pre-treatment concentration. **C:** Time-kill curves after 30 min GEN pre-treatment followed by CAR main treatment. Only inhibitory concentrations of GEN pre-treatment (> MIC) increased killing rates in CAR main treatment. **D-I:** Hysteresis time-kill experiments with three clinically relevant β-lactams (PIT: Piperacillin + Tazobactam; MER: Meropenem; CTZ: Ceftazidime) and gentamicin. For each antibiotic pair hysteresis effects were assessed testing both directions (β-lactam first vs. gentamicin first) using varying concentrations of the pre-treatment drug and two concentrations of the main treatment drug. For β-lactams followed by GEN, negative hysteresis increased with higher pre-treatment concentrations. At higher main treatment concentrations killing saturated. GEN pre-treatment only induced increased killing during β-lactam treatment at concentrations above MIC. Doubling the β-lactam concentration did not result in saturation of the killing effect. The curves are obtained from a single well-growing culture. The raw data is provided in SI Datasets, **Fig. S1**.

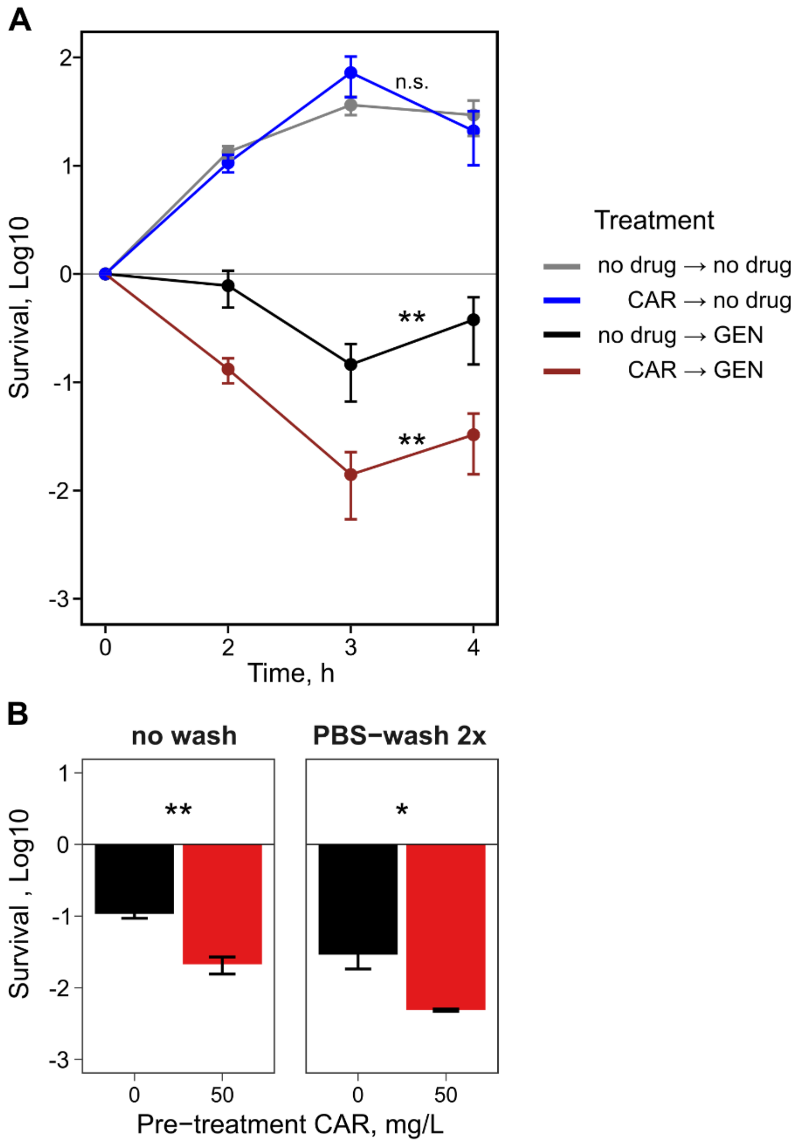


**Fig. S2: Negative hysteresis is distinct from the post-antibiotic effect and is not explained by retention of unbound antibiotic. A:** Pre-treated cells (50 mg/L CAR) grew just as well as untreated cells when resuspended to drug-free medium instead of 1 mg/L GEN (n = 2-4, mean ± SEM); Student's *t-test* adjusted for multiple comparisons using the false discovery rate. With post-antibiotic effect the blue and grey lines would differ significantly. **B**: Hysteresis is not explained by carry-over of unbound pre-treatment antibiotic. Cells are pre-treated for 30 min with a sub-lethal dose of 50 mg/L CAR. Cells are collected by centrifugation, supernatant fully discarded and resuspended for main treatment in GEN 1mg/L and incubated at 37°C ("no wash"). To assess the impact of possible carry-over of unbound pre-treatment antibiotic, cells are alternatively washed twice with antibiotic-free phosphate buffered saline, prior to the begin of main treatment. Survival is assessed by CFU/mL after 4h of GEN treatment. Washes generally reduced survival, but did not affect induction of the hysteresis phenotype, which remained significantly different from the control (n = 3 per treatment, mean ± SEM; Student's *t*-test: *: $P < 0.05$, **: $P < 0.01$). The raw data is provided in SI Datasets, **Fig. S2**.

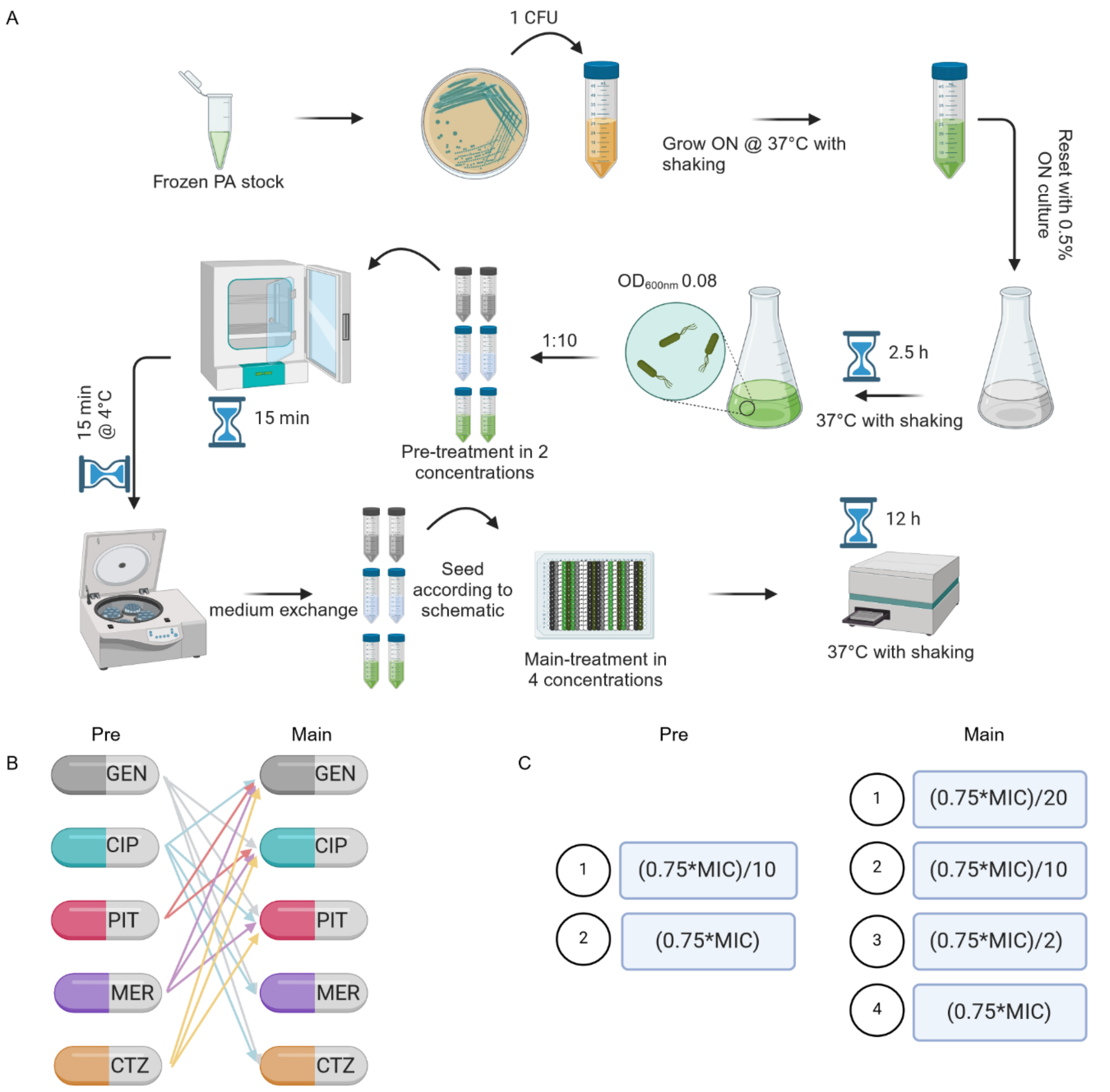


**Fig. S3: Schematics for the hysteresis screen. A:** Experimental procedure. **B:** Antibiotic sequences tested. **C:** Pre and main treatment concentrations tested.

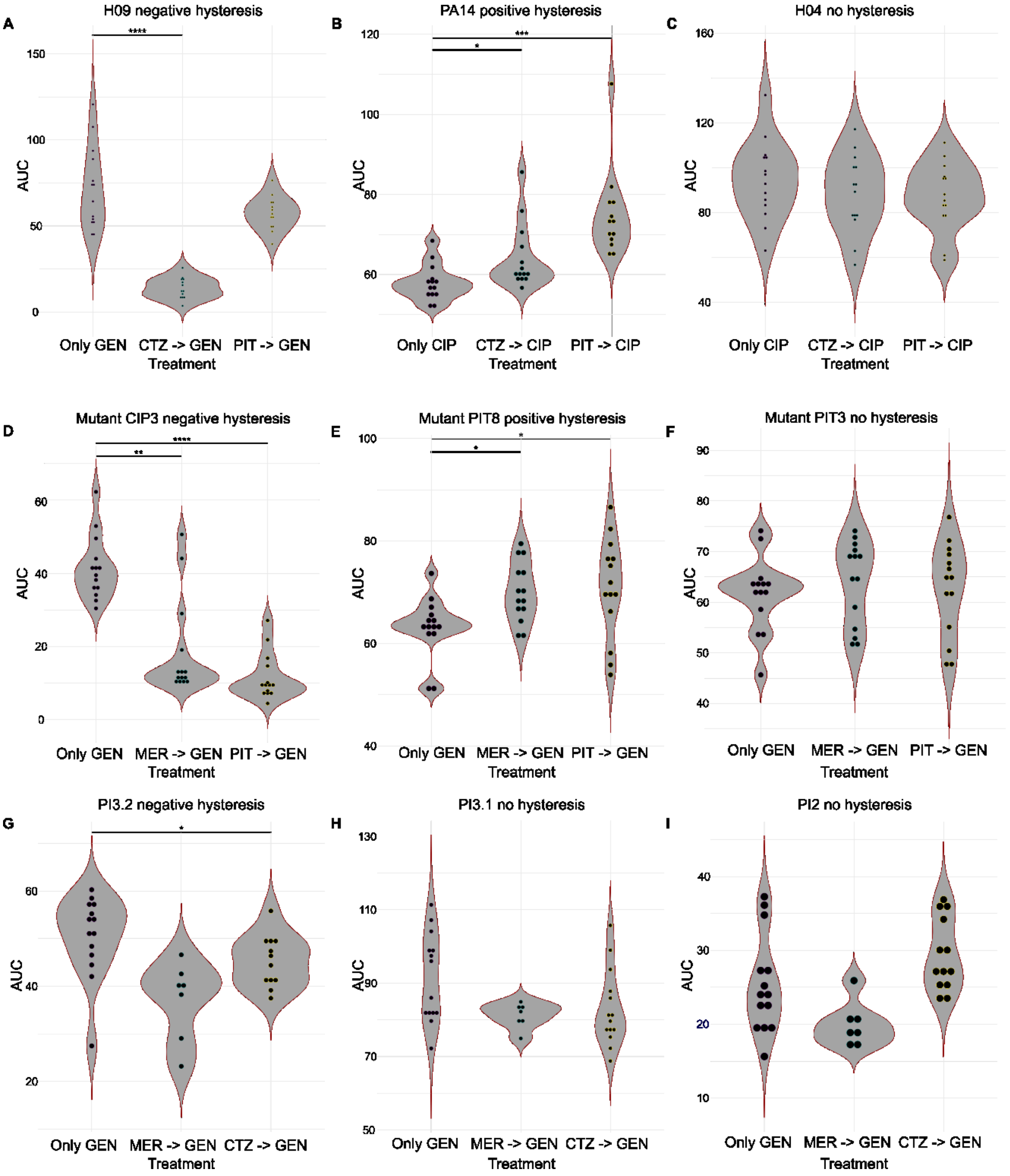


**Fig. S4: Illustrative comparisons of hysteresis types in Fig. 3 A – C:** Example graphs for the mPact panel: **A:** Negative hysteresis. **B:** Positive hysteresis. **C:** No hysteresis. **D – F:** Example graphs for resistant clones: **D:** Negative hysteresis. **E:** Positive hysteresis. **F:** No hysteresis. **G – I:** Example graphs for the COPD within-patient populations: **G:** Negative hysteresis. **H, I:** No hysteresis. AUC (area under the curve) values were visualized as violin plots, with each dot representing a technical replicate. Statistical significance was determined for comparisons of antibiotic switches with the control that only included the main treatment antibiotic (i.e., without pre-treatment), using a Wilcoxon rank sum test, with *P*-values adjusted for multiple comparisons using the false discovery rate (*: $P < 0.05$; **: $P < 0.01$; ***: $P < 0.001$; ****: $P < 0.0001$; n = 7 - 14). Abbreviations: meropenem (MER), ceftazidime (CAZ), piperacillin/tazobactam (PIT), gentamicin (GEN), and ciprofloxacin (CIP). The raw data is provided in SI Datasets, **Fig. S4;** SI Tables**, Table S3**.

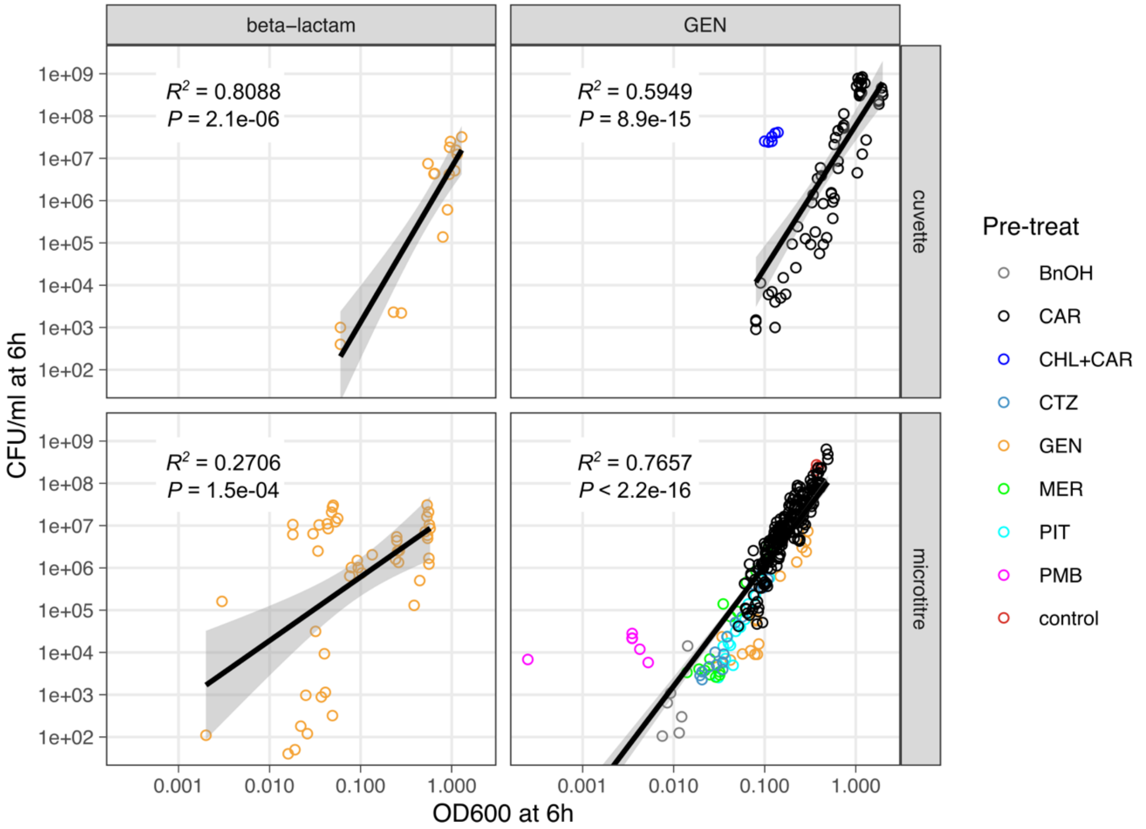


**Fig. S5: Relationship of OD to CFU in hysteresis experiments with *P. aeruginosa* PA14 for paired data points.** Cell population survival after 6 h of main treatment in hysteresis experiments is assessed by optical density at 600 nm (measured in cuvettes or microtiter plates) and by CFU/mL on agar plates. OD and CFU/mL are strongly correlated for GEN main treatments (left columns) across a diversity of pre-treatments. $R^2$ values and $P$ values obtained from Pearson's product moment correlation are specified in each panel. When the CHL+CAR pretreatments are removed, the $R^2$ value for cuvette measurements and GEN main treatment increases to 0.8768. The relationship is less strong when β-lactams are used as main treatment, consistent with their effect on cell morphology. The raw data is provided in SI Datasets, **Fig. S5**.

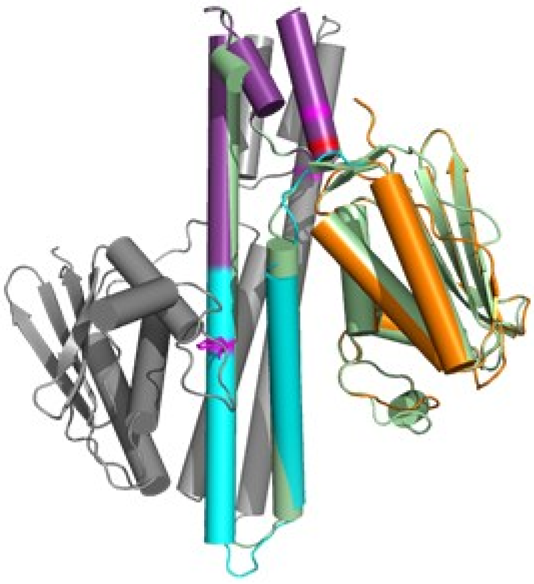

**Fig. S6: Superimposition of the CpxS_HAMP-DHp-CA domain onto CpxA**. CpxA (PDB entry 4BIV) is depicted in green.

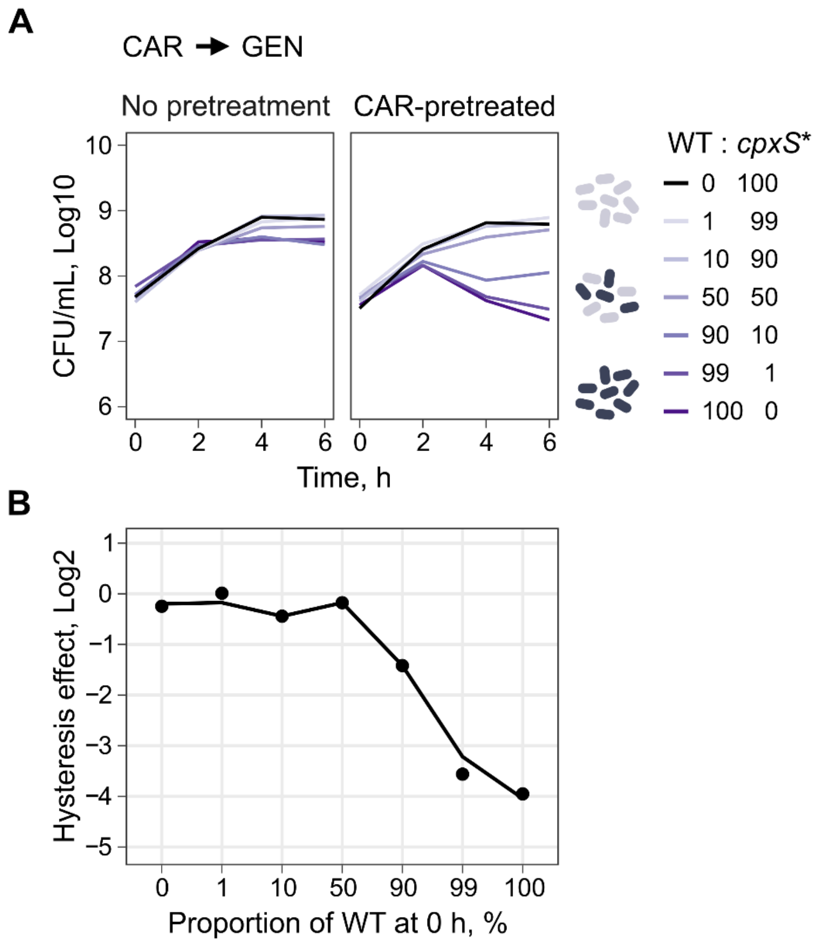


**Fig. S7: CAR → GEN hysteresis experiment using different ratios of PA14 (WT) and CpxS T163P (*cpxS**). A:** Time-kill curves for different mixtures of PA14 and CpxS T163P using 50 mg/L CAR as pre-treatment and 1 mg/L GEN as main treatment. **B:** Summary of the hysteresis effect comparing the no pre-treatment curve with the respective CAR pre-treated curve. The hysteresis effect decreased the higher the proportion of CpxS T163P was in the mixture. At a ratio of 50:50 for both strains there was no longer negative hysteresis. (CAR: Carbenicillin; GEN: Gentamicin). The raw data is provided in SI Datasets, **Fig. S7**.

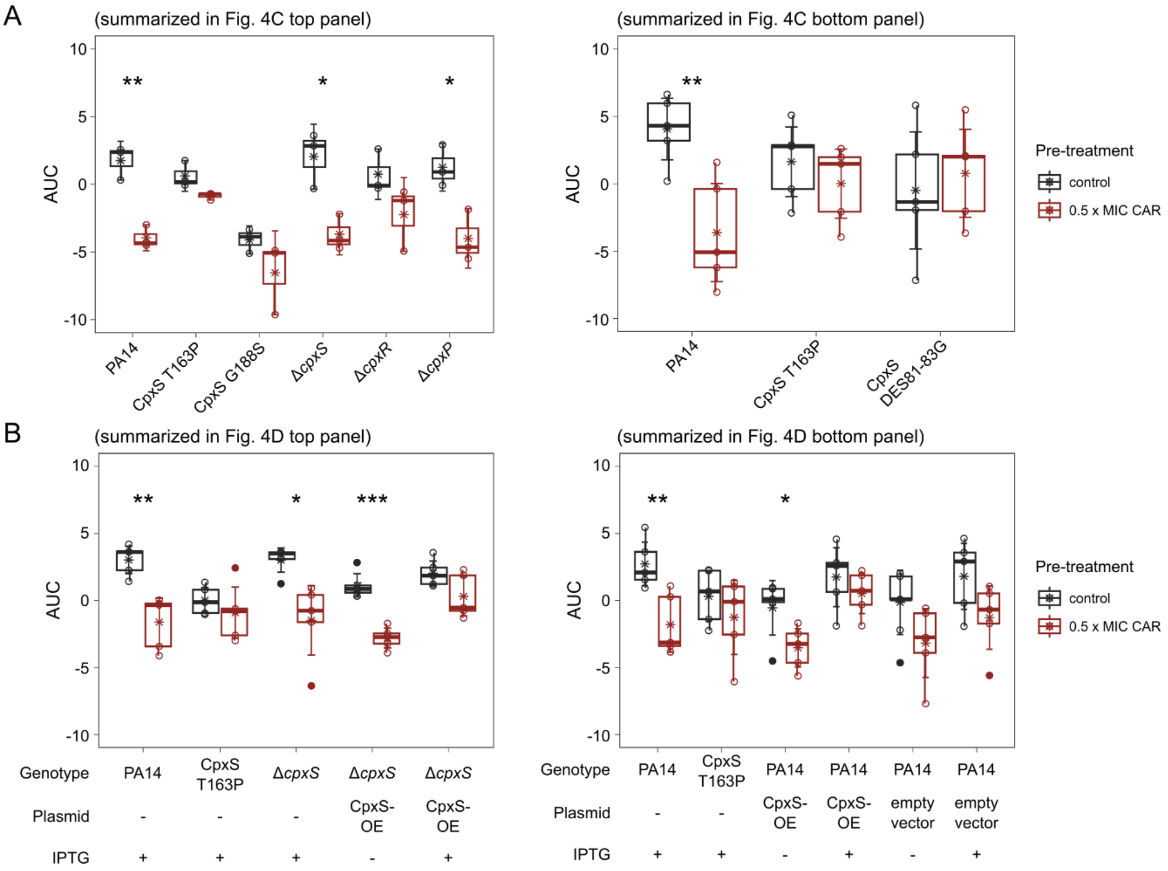


**Fig. S8: Boxplots of AUC data used for statistical analysis of the hysteresis time-kill experiments shown in Figure 4C & D.** For each strain the AUC (area under the curve) data of the control and the pre-treated bacteria are shown as a boxplot. Circles depict individual data points, asterisks indicate the mean, error bars depict mean ± 2 x SEM (standard error of the mean). Student's *t*-test was used to test for presence of negative hysteresis (*: $P < 0.05$; **: $P < 0.01$; ***: $P < 0.001$; n = 3 – 5) comparing both treatments. The raw data is provided in SI Datasets, **Fig. S8**; SI Tables, **Table S8**.

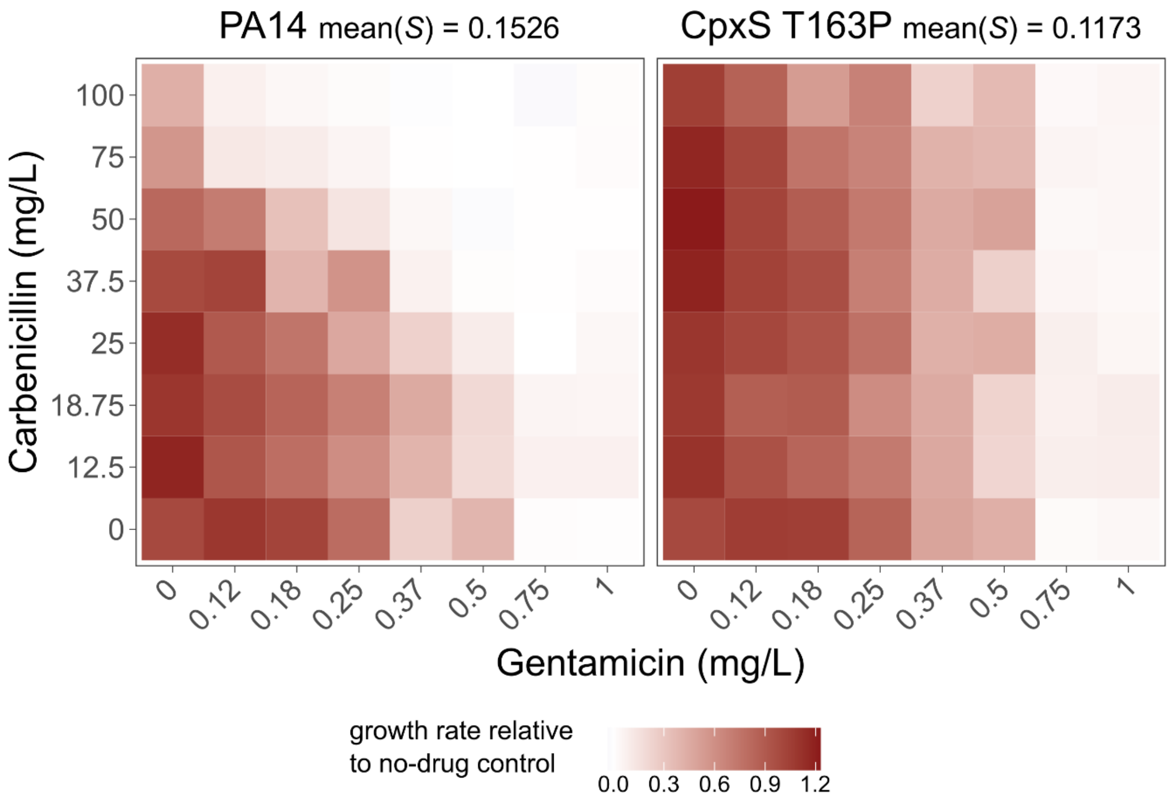


**Fig. S9: CAR – GEN interaction profile for PA14 and CpxS T163P.** The interaction profile was generated using the checkerboard approach. The heatmap shows the mean growth rate for each of the tested drug concentrations normalized by the no-drug control (n = 2). The degree of synergy (*S*) was calculated using the Bliss independence approach. The depicted values summarize the degree of synergy across the whole panel. Positive values depict synergy, while negative values represent antagonism. The raw data is provided in SI Datasets, **Fig. S9**.

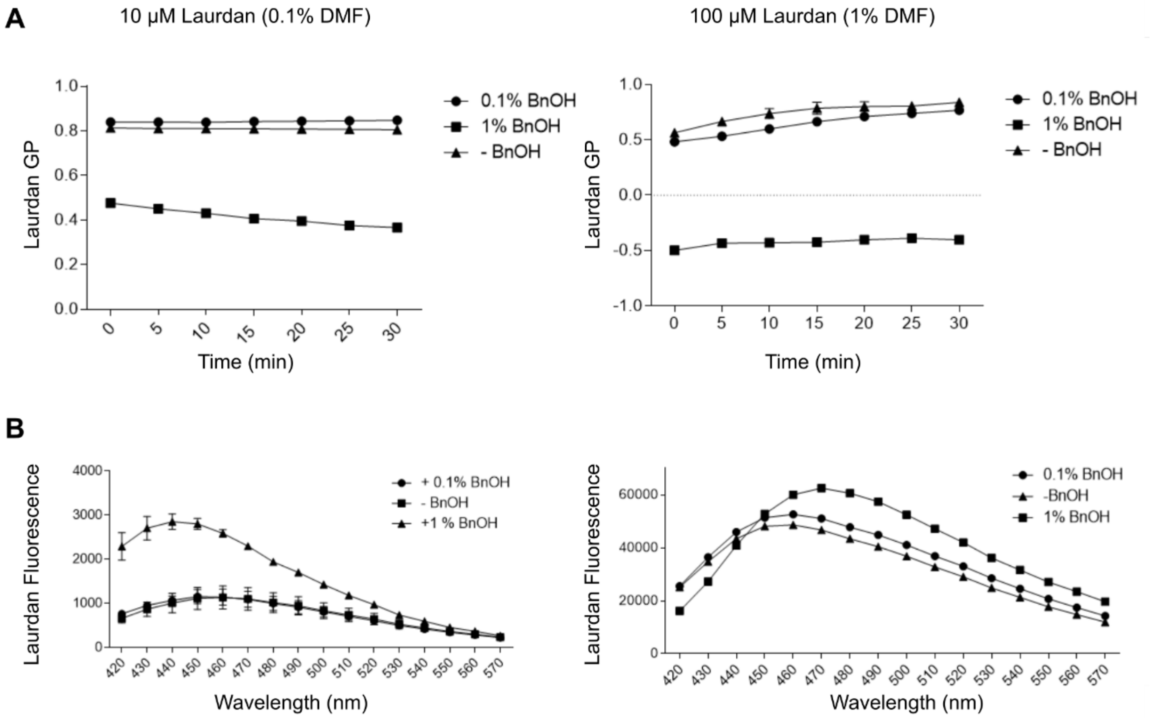


**Fig. S10: Dosage finding of the optimal Laurdan concentration for Gram-negative *P. aeruginosa* cells.** **A:** 10 µM Laurdan, **B:** 100 µM Laurdan. (SI Datasets; **Fig. S10**).

### Extended Methods

#### 1. *Bacterial strains, culture conditions, and antibiotics*

An overview of all bacterial strains, plasmids, and antibiotics used or generated in this project is provided in the **SI Key Resource Table**. Bacterial strains were grown on agar plates or in liquid culture (shaking at 150 rpm) at 37 °C either in M9 minimal medium supplemented with casamino acids (1 g/L), citrate (0.58 g/L), and glucose (2 g/L) or LB-medium. LB-agar was used for counting colony forming units (CFU) and streak-plating of frozen stocks. The strains were stored at -80 °C and stabilized with either 10 % DMSO or glycerol. All media and buffers were autoclaved for sterility, except for antibiotic stocks. Piperacillin was used in combination with the β-lactamase inhibitor tazobactam at a ratio of 8:1 (with the exception of the time-kill experiments for Fig.S3 where 4 mg/L tazobactam were used as assay concentration). All antibiotics were prepared according to the manufacturers' instructions.

#### 2. *Detailed protocol for isolation of clinical isolates*

Clinical Isolates were collected from sputum cultures of patients enrolled in the clinical observational trial "Airway colonization with *Pseudomonas aeruginosa* in chronic obstructive pulmonary disease (COPD)/non-CF bronchiectasis - an observational and biomaterial study" at LungenClinic Großhansdorf (Großhansdorf, Germany). The trial was approved by the local ethics committee of the Medical Faculty of the University of Lübeck (No. 20-295). Further details can be found in the German Clinical Trials Register (https://drks.de/, ID DRKS00023975). Prior to enrollment, patients provided written informed consent for participation. In short, patients with known colonization of the airways by *P. aeruginosa*, admitted to hospital care for acute exacerbation or elective treatment of COPD and/or non-CF bronchiectasis, were enrolled in the trial. Over the course of in-patient treatment as well as at post-discharge follow-up visits, they provided sputum samples for explorative analysis. If bronchoscopy was performed in routine clinical care, bronchial aspirates were additionally collected. Sputum samples and bronchial aspirates were processed and homogenized by incubation with dithiothreitol (Sputolysin® Reagent, Merck; prepared according to manufacturer's instructions). Aliquots for bacterial culture were sent to the laboratory of the Clinic of Infectious Diseases and Microbiology, University Hospital Schleswig-Holstein Campus Lübeck, University of Lübeck, Germany. Bacterial culture was performed according to clinical standard operating procedures. *P. aeruginosa* isolates were harvested, grown in pure culture, and stored at -80 °C until further use. We used six isolate collections from four patients, two patients had two sampling time points, all of which included 8-14 isolates. During bacterial isolation, only three samples had additional bacterial species present (Supplementary **Table S21**, whereby we here focused solely on *P.aeruginosa* as the model pathogen of our study). Further information on the individual isolates, such as morphology and MIC values of the patients, can be found in Supplementary **Table S22**.

#### 3. *Detailed protocols for antimicrobial susceptibility testing*

We used three different methods to measure MIC, depending on the required precision of the MICs for the following assay. The time-kill assays require the most precise determination of MIC values, in order to hit the desired inhibitory concentration; therefore, MICs were determined with the broth micro-dilution approach (BMD). For the large screen, we used either the Vitek2 approach or E-tests to obtain reference MICs and then tested two different pre-treatment concentrations and four different main treatment concentrations. Within the screen, we always selected the treatment that induced the highest amount of inhibition (up to 75%) for our analysis. The MIC determination used was consistent within each panel and assay, thereby ensuring consistency and comparability within each of the performed assays.

#### 3.1 MIC determination with broth microdilution

Determination of MICs with broth microdilution (BMD) was based on the EUCAST (European Committee for Antimicrobial Susceptibility Testing) guidelines for antimicrobial susceptibility testing (1). In a round-bottom 96-well plate a 2-fold serial dilution of the antibiotic in M9 (plus supplements, if indicated) was prepared. 90 µL of diluted antibiotic solution were inoculated with 90 µL of a 1:500 dilution of a 1 mL overnight culture (final inoculum: 1:1000 dilution). The plate was incubated statically at 37 °C for approximately 20 h. After resuspension by pipetting the optical density at 600 nm ($OD_{600}$) was measured in an *EON* Microplate Spectrophotometer (Agilent). MIC was determined as the lowest concentration that results in a relative OD (OD relative to growth control) below 0.1.

#### 3.2 MIC determination with the Vitek®2

For the hysteresis screen, MIC was determined with the Vitek®2 (bioMérieux) diagnostic system according to the manufacturer's instructions. MICs for the mPact panel were previously determined in (2) (SI Tables**, Table S3**).

#### 3.3 MIC determination with MIC test strip

We determined MIC with a MIC test strip (Liofilchem®) for the COPD patient populations. Each isolate was grown individually in 10 mL of LB medium overnight, diluted to $OD_{600}$ ~ 0.08, and mixed at a 1:1 ratio for each patient, respectively. The COPD patient population was then spread onto M9 plates with a cotton swab and incubated for 24 h. MIC was then determined according to the manufacturer's guidelines (SI Tables**, Table S3**).

### 4. *Detailed protocols hysteresis experiments*

#### 4.1 Detailed hysteresis time-kill assay standard protocol (Figure 1 and Figure 4)

To generate exponential phase cells, an overnight culture was diluted 1:200 in M9 (plus supplements if indicated) and incubated at 37 °C until an $OD_{600}$ of 0.06 – 0.07 was reached. At this point, the culture was split into 5 mL portions and the pre-treatment drug was added. Mock treatments without pre-treatment served as control. After incubation at 37 °C for the duration of the pre-treatment (usually 30 min, if not indicated otherwise), the pre-treatment antibiotic was removed by centrifugation of the cultures (10 min, 4 °C, 3220 x g). The supernatant was discarded and the pellet was resuspended in 5 mL fresh medium (RT). Survival of cells was evaluated by colony forming units (CFU) per mL culture. Initial CFU density was determined by taking a 20 µL sample following culture resuspension ($t_0$). The main treatment drug was immediately added, and the samples were incubated at 37 °C. After 2, 4, and 6 h of incubation, additional 20 µL samples for CFU determination were taken. For CFU determination, a 10-fold dilution series in 0.9 % sterile-filtered NaCl was performed. In assays with overexpression mutants, phosphate buffered saline (PBS) supplemented with $CaCl_2$ and $MgSO_4$ (final concentrations 0.1 mM and 1 mM, respectively) was used instead of 0.9 % NaCl in order to minimize potential killing during dilution (3). 100 µL of diluted samples were plated on LB agar plates and incubated overnight at 37 °C. The next day, CFU were counted.

In assays with overexpression vectors, IPTG was also added to the controls in order to exclude possible side effects of IPTG.

Time-kill curves were usually plotted as survival over time. Survival was calculated based on the obtained CFU values using the following equation:

$$Survival = \frac{CFU}{CFU_{t_0}}$$

The area under the curve (AUC) of the $\log_{10}$-transformed survival data was calculated for each treatment and compared using Student's *t*-test for each strain.

**4.2 Detailed protocol for hysteresis time-kill experiments with clinically relevant antibiotics (Figure 2A)**

To generate exponential phase cells, an LB overnight culture was diluted 1:200 in M9 and incubated at 37 °C until an $OD_{600}$ of 0.08 was reached. Then 1 mL of exponential phase cultures was added to 9 mL of pre-warmed M9 medium in a 50 mL tube to avoid the inoculum effect (4) and the pre-treatment drug was added (where appropriate). After incubating the tubes at 37 °C for 15 min, the samples were centrifuged for 15 min (4 °C, 3220 x g). The supernatant was discarded and the pellet was resuspended in 10 mL pre-warmed M9 medium. Initial CFU density was determined by taking three 20 µL samples from each tube following culture resuspension ($t_0$). Next, the main treatment antibiotic was added and the tubes were incubated at 37 °C for 6 h. After 2, 4, and 6 h of incubation, additional samples for CFU determination were taken. For CFU determination, a 10-fold dilution series in PBS was performed. 7 µL of diluted samples were spotted on LB agar plates and incubated overnight at 37 °C. The next day, CFU were counted. The area under the curve (AUC) of the obtained time-kill curves was calculated based on the $log_{10}$-transformed CFU data. The hysteresis effect was quantified as ΔAUC (hysteresis treatment – same pre-treatment control) and summarized in a heatmap.

<u>Determination of drug concentrations to be used in hysteresis time-kill experiments:</u>
The concentrations to be used in hysteresis time-kill assays were determined using the same experimental set up as for a regular hysteresis experiment. During the pre-treatment period all samples were incubated without any antibiotic. During the main treatment period multiple concentrations of the antibiotic of interest were tested. CFUs were determined at the beginning of the main treatment (before the antibiotic was added) and after 6 h of incubation. A dose-response-curve was plotted for the t = 6 h data and concentration that reduced cell density by two orders of magnitude was determined by manually inspecting the curves. The chosen doses for each antibiotic were tested in a confirmation run, before performing the actual hysteresis time-kill experiments.

**4.3 Pause assay (Figure 1*G*)**

To elucidate how a growth period with no drug effects hysteresis (i.e. introducing a pause), we adapted our standard CFU-based hysteresis protocol (described above) by introducing drug-free growth periods. In short, all cells were pre-treated with carbenicillin for 30 min and spun down for 10 min (3220 x g, 4 °C). The medium was exchanged for all samples with drug-free medium and the samples were incubated at 37 °C shaking at 150 rpm. Each sample received GEN main treatment after 0, 10, 20, 40 or 80 min of growth in drug-free medium. All samples with drug-free growth periods were centrifugated again (10 min, 3220 x g, 4 °C), received new media and got their OD adjusted to 0.08 to control for cell density. Sampling was similar to the standard hysteresis protocol after 0, 2, 4, and 6 h of GEN treatment. CFUs were determined by spotting 7 µL of a 10-fold dilution series in PBS onto LB-agar plates. After incubating for 20 h at 37 °C the CFU/mL was determined.

AUC of the time-kill curves was calculated using the $log_{10}$-transformed CFU data. The AUC data were then used to generate a Nelder-Mead dose-response model (5) of the hysteresis effect size depending on the duration of the pause.

**4.4 Hysteresis screen experimental design and protocol (Figure 3)**

The rationale of the hysteresis screen was to provide a high-throughput screen that reduces preliminary experiments such as MIC determination via broth microdilution. We determined the MIC with the clinical diagnostics machine Vitek®2 (bioMérieux). We here used five drugs regularly used for *P. aeruginosa* infections with different functional targets. We used piperacillin/tazobactam (PIT), ceftazidime (CTZ), meropenem (MER), ciprofloxacin (CIP), and gentamicin (GEN). Especially for some antibiotics, e.g., CIP and MER, most of our strains had a MIC below the detection limit (SI Tables**, Table S3**). Therefore, we chose to test two pre-treatment concentrations, 0.75 x MIC and (0.75/10) x MIC. Similarly, we also tested four different main treatment concentrations, namely,0.75 x MIC, (0.75/2) x MIC, (0.75/10) x MIC, and (0.75/20) x MIC. We chose these concentrations with the intention to achieve concentrations in which (i) no effect of the pre-treatment

is present and (ii) partial inhibition of the main treatment concentrations is achieved. All antibiotic classes were reciprocally tested against each other, and we additionally included two intra-class comparisons, MER/CTZ → PIT. We used 384-well plates (Greiner) to increase throughput, which allowed us to test two pre-treatment antibiotics for one main treatment antibiotic. We randomized the columns to prevent gradient effects within the plate reader. The screen was designed to provide a binary yes or no answer and not to determine hysteresis effect size. All bacteria were streaked on an LB agar plate and incubated for 24 h. One CFU was then used to inoculate 10 mL of LB broth. 0.5 % overnight culture was subsequently reset in 100 mL of fresh M9 media and incubated until an $OD_{600}$ of 0.075 – 0.08 was reached, corresponding to exponential phase cells (5 x $10^7$ CFU/mL). To avoid the inoculum effect (4), the exponential phase culture was diluted 1:10 in 15 mL M9 containing the corresponding treatment. The cells were incubated in the pre-treatment for 15 min, followed by 15 min centrifugation (3220 x g at 4 °C). The M9 was discarded, and the cells were resuspended in 7.5 mL of fresh, pre-warmed (37 °C) M9. Then, 50 µL of cells were seeded in 384-well plates that contained either 50 µL of plain medium or main-treatment antibiotics. The plates were incubated at 37 °C with continuous shaking (Shake mode: double orbital; Orbital frequency: 807 cpm (1 mm); Orbital speed: Fast), and $OD_{600}$ in a plate reader (Epoch 2, Agilent) for 12 h in 15 min intervals. An overview of the experimental set up is provided in **Fig. S4**.
We would like to emphasize that this assay was developed as a screening tool and, therefore may be unable to optimally assess individual strains as the effective concentration used may vary between strains. The initial MIC was determined with automated susceptibility testing optimized for detecting clinical resistance breakpoints. Such automated testing is inherently less precise than determining MIC with broth micro dilutions where antibiotic concentrations, media, and other experimental conditions can be changed (6, 7). For example, in strains H01 and H03, we detected positive hysteresis in β-lactam → GEN treatments, both strains are of clinical origin and are highly resistant to the used antibiotics (2). For these strains, we only achieved 10 % growth inhibition. Additionally, β-lactams cause filamentation, which inflates the optical density (8). These factors make our assay prone to falsely identifying positive hysteresis. Despite these limitations, we deem our OD-based methodology appropriate for the primary focus of our study: the investigation of the distribution of negative hysteresis across bacterial strains.

#### 4.5 Hysteresis screen data and statistical analysis

Raw data from the plate readers were quality controlled within an R-based pipeline, which entailed the following parts. (i) De-randomization. (ii) Removal of rows and wells: We discarded the top and bottom rows, accounting for potential error due to evaporation, and additional wells in case of pipetting error. (iii) Spike artifact correction: Spikes in OD values were identified per well by comparing each value with the mean of the two neighboring values. We defined a cut-off of 0.05 to detect most spikes while minimizing false positives effectively. (iv) Outlier Detection and Removal: Outlier OD values were identified by comparing each value against identical treatment replicates for a given time point. Outliers were flagged using a z-value > 3 and a distance of more than 3 interquartile ranges from the first and third quartile and omitted if both conditions were true. After these initial steps, our data contained roughly 3.5 million observations, which we subsequently prepared for analysis. For this, we removed any values containing NAs and changed negative OD values to 0. Additionally, we removed all samples that did not include the first and last time points, as equal length is essential when comparing the area under the curve (AUC). AUC was determined for each tested well with the R package *DescTools* using the spline method. Overall, we evaluated the growth kinetics of around 68,000 wells across the screen. Given the two different pre-treatment concentrations per antibiotic, we had to select one for further analysis. One assumption of the hysteresis assay is that the pre-treatment does not inhibit growth. Therefore, we checked for inhibition with a one-sided Wilcoxon rank sum test. We did not adjust for multiple comparisons to keep the value more stringent. In the case of inhibition, the lower pre-treatment concentration was chosen. In cases where both pre-treatments induced an effect, we chose the lower pre-treatment but excluded these strains from further statistical analyses. The main treatment concentration was selected based on the highest inhibition (max. 75 %) from the four concentrations. Hysteresis presence was then assessed with pair-wise Wilcoxon rank sum tests, comparing the AUC of the hysteresis treatment against the AUC of the main treatment control. *P*-values were corrected for multiple testing using false discovery rate. To avoid bias by including samples in which the pre-treatment had an inhibitory effect, we excluded any significant values we got from them and only depicted the directionality.

### 5. *Detailed protocols drug interactions*

#### 5.1 CombiANT

CombiANT assays (9–11) were performed using M9 agar prepared with Bacto™ Agar (BD, order ID 214010). The inserts were loaded with 500 µL of the respective antibiotic diluted in agar (at 65 °C) and let solidify in the fridge for at least 1 h. To activate the plates 25 mL of agar were pipetted into the center of the insert. The plates solidified for 2-3 h under light protection. Then a 1:50 dilution of an overnight culture in 0.9 % NaCl was used to inoculate the plate by densely streaking with a sterile cotton swab in three directions. After 24 h of incubation at 37 °C, a picture of the plate was taken with the BIO-RAD ChemiDoc Touch (Colorimetric blot, Epi white, medium size (15.5 x 12.4 cm), auto optimal exposure). The obtained images were analyzed using the CombiANT Imager v.3.5 (for Windows) and the nps_v3 analysis pipeline (12).
Comparison of the interaction profile between strains was performed by Wilcoxon rank sum test of the FICi values for each antibiotic pair separately.

#### 5.2 Checkerboard assay

A checkerboard approach was performed to measure the type of interaction between the antibiotics CAR and GEN for the WT strain PA14 and the CpxS T163P mutant. Seven concentrations of each antibiotic and three no-drug controls were distributed across a 96-well plate. CAR concentrations were 12.5 – 100 mg/L and GEN concentrations were 0.12 – 1 mg/L. Each strain was tested twice. The plates were incubated for 24 h at 37 °C with constant shaking and $OD_{600}$ measurements taken every 15 min. We calculated the growth rate *r* for each well by fitting a linear regression of growth over time during the exponential phase. Exponential phase was generally observed between 195 to 360 min (13). The growth rates were then normalized to the mean of the no-drug controls for each strain and replicate. In the heatmap (**Figure S9**) the mean normalized growth rate of both replicates is plotted. Subsequently, we determined the degree of synergy using the Bliss independence method described previously (14) based on the mean growth rate of both replicates:

$$S = (r_{A0}/r_{00})(r_{0B}/r_{00}) - (r_{AB}/r_{00})$$

where $r_{A0}$ represents the growth rate at a given concentration of drug A in the absence of B and vice versa for $r_{0B}$. $r_{00}$ is the growth rate of the no-drug control and $r_{AB}$ is the growth rate at any concentrations of the drugs A and B being combined. The degree of synergy *S* was only calculated for drug combination with growth rate ≥ 0. Positive values indicate synergy, while negative values indicate antagonism.

### 6. *Detailed transcriptomics*

Cells of PA14 and Cpx T163P were grown to mid-exponential phase ($OD_{600}$ = 0.065) at 37 °C, as described in the standard hysteresis time-kill protocol, and then treated with 50 mg/L CAR for hysteresis induction or not treated. After 30 min, cells were resuspended in antibiotic-free medium and harvested at three sequential time points (0, 30 and 60 min) to investigate the expression dynamics induced by short-term CAR pretreatment over time. Cells were collected by centrifugation (3220 x g, 10 min, 4 °C) and total RNA was isolated using a TRIzol lysis and the NucleoSpin miRNA kit (Macherey-Nagel). RNA was eluted in TE buffer and stored at -80 °C prior to rRNA depletion (RiboZero Bacterial Kit), library preparation (TruSeq stranded total RNA) and sequencing (Illumina, HiSeq4000, 1 x 50 bp). In silico quality control was carried out via Fast QC (v0.11.9) and MultiQC, part of Miniconda3 (v4.8.2). Reads were trimmed with Trimmomatic (15) v0.39, settings phredThreshold=phred33, seedMismatches=2, palindromeClipThreshold=10, lengthMin=36). The reference genome for UCBPP-PA14 was retrieved from GenBank (NCBI Reference Sequence NC_008463.1) and for the Cpx T163P mutant, the genome was adapted via a custom R script to include the mutation. Reads were followingly mapped to the respective genomes using EDGEpro (16) (v1.3.1), substituting the bundled Bowtie package (17) with the newest available at the time of mapping (Bowtie2 v2.4.1). Gene, Gene Ontology and KEGG pathway annotations were downloaded from pseudomonas.com (database version 19.1). Read count normalization and statistical analysis was carried out in R (v3.6.1) via edgeR (18) (v3.28.1), enrichment analyses were performed with clusterprofiler (v3.14.3) Differential expression between CAR-induced and uninduced cells was determined, focusing on time point 0 and thus at the time when negative hysteresis is most strongly expressed and before it ceases. We also assessed differential expression between the PA14 wildtype and the CpxS T163P mutant in the absence of a pre-treatment across the three considered time points.

### 7. *Detailed membrane stress measurements*

The *P. aeruginosa* strain PA14 and the mutant CpxS T163P were grown overnight at 37 °C in M9 medium, subsequently diluted by 0.05 % in fresh media, and grown to logarithmic growth phase ($OD_{600}$ = 0.1). Cells were pre-treated with CAR (50 mg/L) and after 15 min Laurdan (100 µM, ThermoFischer Scientific) was added. Optimal Laurdan concentration was evaluated before to ensure proper staining of the Gram-negative *P. aeruginosa* cells, as the original protocol used was designed for Gram-positive cells (19, 20) (**Figure S10**). Stained and pre-treated cells were collected after 30 min by centrifugation at 16,000 x g for a minute at 37 °C, washed twice in M9 containing 0.8 % DMF, and finally resuspended in M9. Cells were transferred into a black polystyrene 96-well flat bottom plate (ThermoFisher Scientific). All measurements were conducted in an Infinite M Plex plate reader (excitation 350 nm, emission 455 nm and 505 nm, Tecan). Laurdan General Polarization (GP) was calculated from the spectra with the following equation:

$$GP = \frac{I\,(455nm) - I\,(505nm)}{I\,(455nm) + I\,(505nm)}$$

where *I* denotes the fluorescent intensity. The impact of CAR on membrane fluidity was assessed by comparing CAR pre-treated bacteria to no drug controls (*t*-test).

### 8. *Detailed determination of intracellular GEN concentrations with ELISA*

The overall experimental setup was similar to a standard hysteresis assay (see above). In order to allow for sampling of higher volumes, we increased the culture volume to 6 mL (instead of the usual 5 mL). 1 mL samples for GEN concentration measurement were taken 20 and 40 min after GEN addition. In parallel, 20 µL samples (3 technical replicates) for CFU determination were taken. Both strains (PA14 and CpxS T163P) were tested in biological triplicates.

For CFU determination, a 10-fold dilution series in PBS was performed. 7 µL of diluted samples were spotted on LB agar plates and incubated overnight at 37 °C. The next day, CFU were counted.

The samples for GEN concentration measurement were pipetted into a DNA low-bind tube and centrifugated (5 min, max. speed, 4 °C). The supernatant was discarded and the cells were washed 3 times with 0.5 mL PBS. After the last washing step, the cells were resuspended in 250 µL lysis buffer (1x TE buffer (10 mM TRIS, 1 mM EDTA, pH 8) supplemented with 10 mg/mL lysozyme (Carl Roth, 8259.1) and 2.2 mg/mL proteinase K (Carl Roth, 7525.2)) and incubated for 30 min at 37 °C. Then the cell debris was spun down (10 min, max. speed, RT) and the supernatant was diluted 1:5 in lysis buffer. Next, the GEN ELISA (Elabscience®, E-FS-E073) was performed in technical duplicates according to the manufacturer's protocol.

GEN concentrations were calculated based on the obtained standard curve following the manufacturer's instructions. The CFU and GEN concentration data of the technical replicates were summarized as mean. The summarized data were then used normalize the data as GEN concentration/CFU to account for differences in cell density. For statistical analysis, ANOVA of the normalized data was performed for each strain separately (SI Tables, **Table S19**), followed by Tukey HSD post-hoc testing with FDR correction for multiple comparisons (SI Tables, **Table S20**).

### 9. ***Detailed protocol for the construction of mutants by two-step allelic exchange***

We used a previously established two-step allelic exchange approach (21, 22) to generate scar-free deletion or SNP (single nucleotide polymorphism) mutants of *P. aeruginosa* genes (for simplicity, SNP mutants include mutants with small deletions (up to 10 bp) in this study). Briefly, fragments of approximately 700 bp up – and downstream of the mutation of interest were cloned into the plasmid pUIsacB (22), containing a tetracycline (TET) resistance (*tetA*) and a sucrose sensitivity (*sacB*) gene, using Gibson Assembly. The resulting plasmid was transformed into *E. coli* JM109. In a tri-parental conjugation, the construct was inserted into the chromosome of *P. aeruginosa* by homologous recombination. This step resulted in merodiploid bacteria that were tetracycline resistant and sucrose sensitive. A second selection step on sucrose medium selected for bacteria that had looped out the *tetA* and *sacB* gene by homologous recombination resulting in bacteria with either the wild-type or the mutated allele at an expected ratio of 50 % each.

#### 9.1 Primer design

Primers were designed based on the PA14 reference genome (NCBI: NC_008463.1). To amplify fragments for Gibson Assembly from PA14 (or evolved PA14 isolate) DNA primers were designed using *NEBuilder*. Fragments for the generation of deletion mutants were designed to have a length of approximately 700 bp flanking the gene of interest up- and downstream. Sequencing primers were designed approximately 50 bp outside of the construct area but also inside the construct to ensure full sequencing coverage across the construct. For the generation of SNP mutants, fragments were designed to have a length of approximately 1400 bp with the mutation of interest located in the middle. Sequencing primers for SNP mutants were designed approximately 450 bp away from the mutation of interest. A list of all primers used for the construction of mutants by two-step allelic exchange is given below. In this study all primers are referred to by their ID.

Primers used for two-step allelic exchange:

| ID | Primer | Sequence 5' → 3' |
|---|---|---|
| SHP 1065 | pUIsacB_F | AGCACTACATCAACTGACTA |
| SHP 1066 | pUIsacB_R | CTGAACCAAGATAGCTGTAC |
| SHP 1067 | pUIsacB-seq_F | AGCGTTCTGAACAAATCCAGATG |
| SHP 1068 | pUIsacB-seq_R | ATTTGTCCTACTCAGGAGAGCGTTC |
| SHP 1087 | cpxS-up_F | gtacagctatcttggttcagCTTGACCTCCCTAGACTGG |
| SHP 1088 | cpxS-up_R | cggtcgccttGGTTTTCCTGTTGAATCGC |
| SHP 1089 | cpxS-down_F | caggaaaaccAAGGCGACCGAATAACGC |
| SHP 1119 | cpxS-down-new_R | tagtcagttgatgtagtgctGCTTCGTCTACCTGGGTAC |
| SHP 1105 | cpxS-seq_F | CCACTGAGCGGGTTTCGTCAG |
| SHP 1120 | cpxS-seq-new_R | GAGATGGCCACCGATCCGGT |
| SHP 1123 | cpxS-seq_I2 | GCCTGGCCGAGGATGTCCAG |
| SHP 1124 | cpxS-seq_I1 | GCGGCCCGATGAAGGAACTG |
| SHP 1135 | cpxS-V2_R [1] | gtcagttgatgtagtgctACTTCCAGGCGGATATCCTG |
| SHP 1136 | cpxS-V2_F | cagctatcttggttcagATGGAAGGCGGCCACATGATG |
| SHP 1141 | cpxS-V2-seq_F | GCAGCAGAAGAAGTTCGACGAA |
| SHP 1142 | cpxS-V2-seq_R | AGTTCGTGGGACACGTCGCGGA |
| SHP 1137 | cpxS-V1_R [2] | gtcagttgatgtagtgctGTTATTCGGTCGCCTTGCGC |
| SHP 1138 | cpxS-V1_F | cagctatcttggttcagACGAACTGCAGAAGAAGCGC |
| SHP 1139 | cpxS-V1-seq_F | CTGAAGGATCTCGCCGAGCAGT |
| SHP 1140 | cpxS-V1-seq_R | TCGAACATGTCCGGCCAGCCGT |
| SHP 1161 | cpxR-up_F | gtacagctatcttggttcagATCGACAGCAACGACCCG |
| SHP 1162 | cpxR-up_R | aacccgctcaCGGGTGTCGCTCAATTGATC |
| SHP 1163 | cpxR-down_F | gcgacacccgTGAGCGGGTTTCGTCAGC |
| SHP 1164 | cpxR-down_R | tagtcagttgatgtagtgctAGAAGGCGGCGAGGATTC |
| SHP 1165 | cpxR-seq_F | ATCGACAGCAACGACCCG |
| SHP 1166 | cpxR-seq_R | AGAAGGCGGCGAGGATTC |
| SHP 1167 | cpxR-seq_I1 | AACCCGCGCCAGCACCGATT |
| SHP 1168 | cpxR-seq_I2 | TTGCCCATCTGCTGGCGCTG |
| SHP 1169 | cpxP-up_F | gtacagctatcttggttcagGGTCCAGGAAGGTTTCTC |
| SHP 1170 | cpxP-up_R | gtcagttggaGGTGTTTCTCCTTTCTGGG |

| | | |
|---|---|---|
| SHP 1171 | cpxP-down_F | gagaaacaccTCCAACTGACGTCCGAGC |
| SHP 1172 | cpxP-down_R | tagtcagttgatgtagtgctTTGAAATCGCGCGCCAGG |
| SHP 1173 | cpxP-seq_F | GGTCCAGGAAGGTTTCTC |
| SHP 1174 | cpxP-seq_R | CCATGCGGTTGAAATCGC |

[1] cpxS-V2: CpxS DES81-83G; [2] cpxS-V1: CpxS G188S.

### 9.2 PCR amplification and assembly of construct components

#### 9.2.1 Amplification and purification of up- and downstream fragments for deletion mutants

Up- and downstream fragments were amplified from genomic PA14 DNA by PCR using Phusion DNA polymerase (Thermo Scientific™; F530S) following the manufacturer's protocol. Genomic DNA was isolated beforehand using the *NucleoSpin Tissue Mini kit for DNA from cells and tissue* (MACHEREY-NAGEL GmbH & Co. KG; 740952.50) according to manufacturer's instruction following the support protocol for bacteria. Primer pairs, annealing temperatures and expected fragment sizes are listed below.

| **Fragment** | **Primer pair** | **$T_A$ (°C)** | **Size (bp)** |
|---|---|---|---|
| cpxS-up | SHP 1087 + SHP 1088 | 60.7 | 645 |
| cpxS-down | SHP 1089 + SHP 1119 | 62.3 | 634 |
| cpxR-up | SHP 1161 + SHP 1162 | 64.1 | 749 |
| cpxR-down | SHP 1163 + SHP 1164 | 64.8 | 753 |
| cpxP-up | SHP 1169 + SHP 1170 | 59.7 | 744 |
| cpxP-down | SHP 1171 + SHP 1172 | 64.8 | 767 |

Fragment sizes were confirmed by agarose gel electrophoresis. The fragments were purified by pooling multiple PCR reactions of the same fragment together followed by gel extraction with the *GeneJET Gel Extraction Kit* (Thermo Scientific™; K0692) according to the manufacturer's manual.

#### 9.2.2 Amplification and purification of fragments for SNP mutants

Genomic DNA isolated from evolved PA14 isolates with *cpxS* mutations (23) (CpxS G188S: isolate 12-1a-D8-18; CpxS DES81-83G: isolate 12-1a-D2-7) was used as template DNA for PCR amplification with Phusion DNA polymerase following the manufacturer's instructions. Primer pairs, annealing temperatures and expected fragment sizes were the following:

| **Fragment** | **Primer pair** | **$T_A$ (°C)** | **Size (bp)** |
|---|---|---|---|
| CpxS G188S | SHP 1137 + SHP 1138 | 66.6 | 1430 |
| CpxS DES_81-83G | SHP 1135 + SHP 1136 | 64.3 | 1530 |

Fragment sizes were confirmed via agarose gel electrophoresis and the fragments were purified by gel extraction as described above.

#### 9.2.3 Amplification and purification of pUIsacB backbone

The pUIsacB plasmid was isolated from *E. coli* Top10 using the *QIAprep Spin Miniprep Ki*t (QIAGEN; 27104) according to manufacturer's instructions. For linearization of the plasmid, 200 ng of the plasmid were digested with *Spe*I-HF (New England Biolabs; R0133S) following the manufacturer's protocol. Afterwards, the linearized plasmid was used to amplify the pUIsacB backbone (~5.2 kb) by PCR with Phusion DNA polymerase (Primers: SHP 1065 & SHP 1066; $T_A$: 62 °C). 5 µL of the PCR product were used to check the size of the amplified product by agarose gel electrophoresis. The remaining 45 µL of PCR product were digested with 1 µL *Dpn*I (New England Biolabs; R0176S) for 1 h at 37 °C in order to remove any template plasmid. Then, the pUIsacB backbone was purified using the *GeneJET PCR Purification Kit* (Thermo Scientific™; K0701) according to the manufacturer's protocol.

### 9.3 Gibson Assembly

Gibson Assembly of the fragments and plasmid backbone was performed using the *NEBuilder® HiFI DNA Assembly Master Mix* (New England Biolabs; E2621L) following the manufacturer's instructions. The molar vector to insert ratio was 1:2 and the assembly reaction was incubated for 1 h at 50 °C.

#### 9.4 Transformation of *E. coli* JM109 and confirmation of transformants

The assembled plasmid was transformed into competent *E. coli* JM109 (Promega) using a heat-shock based approach following the manufacturer's protocol. For transformation 2 µL of assembly product were used. 100 µL of undiluted and 1:10 diluted transformation mix were spread on pre-warmed LB agar + TET 25 mg/L plates. The plates were incubated overnight at 37 °C and screened for GFP (green-fluorescent protein)-positive clones using a fluorescence dissecting microscope the next day. For each construct, multiple clones were picked and the presence of the pUIsacB plasmid was confirmed by colony PCR using DreamTaq DNA polymerase (Thermo Scientific™; EP0701) following the manufacturer's instructions (Primers: SHP 1067 & SHP 1068; $T_A$: 52 °C, initial denaturation: 6 min). The size of the PCR products (~1850 bp or ~250 bp without insert) was checked by agarose gel electrophoresis and cryo-cultures of the picked clones containing the vector with insert were prepared. To confirm correct insertion, PCR products were Sanger sequenced (Primers SHP 1067 and SHP 1068) by the Institute of Clinical Molecular Biology in Kiel or Eurofins Genomics (TubeSeq Service). Samples for sequencing were prepared according to company guidelines. Sequences were analysed with the Basic Local Alignment Search Tool (BLAST) from the National Center for Biotechnology Information, U.S. National Library of Medicine (NCBI) (24, 25).

#### 9.5 Tri-parental conjugation (first crossover) and selection for *P. aeruginosa* transconjugants

In a tri-parental mating the plasmid was transferred from *E. coli* JM109 transformants to PA14. For tri-parental conjugation, overnight cultures of PA14 (recipient), HB101: pRK2013 (helper) and a confirmed JM109 transformant (donor) in LB (+ antibiotic) were prepared. 1 mL of recipient culture was heat-shocked for 20 min at 45 °C. Afterwards, recipient, 0.5 mL donor and 0.5 mL helper cultures were centrifuged for 5 min at 6,000 x g. The supernatants were discarded and all strains were combined in 100 µL LB. Next, the whole conjugation mix was spotted on a LA plate and incubated overnight upright at 37 °C. The next day, the whole lawn was scraped of the plate and resuspended by vortexing in 1 mL LB. 100 µL of resuspended cells (undiluted, 1:10 dilution) were spread on LA + TET 25 mg/L + NF 100 mg/L plates to select for *P. aeruginosa* merodiploids and to kill off the *E. coli* helper and donor strains. Plates were incubated overnight at 37 °C. Merodiploids were identified by their GFP-signal using a fluorescence dissecting microscope.

#### 9.6 Sucrose counter-selection (second crossover) and confirmation of mutants

In a counter-selection step on sucrose media, it was selected for bacteria that had looped out the plasmid. For this purpose, up to 4 merodiploids were picked, streaked on TYS10 plates (10 g/L peptone, 5 g/L yeast extract, 15 g/L Agar-Agar, 10 % w/v sucrose) and incubated overnight at 37 °C. In addition, cryo-cultures of the picked clones were prepared as a back-up. The next day, non-fluorescent colonies were picked from the TYS10 plates and re-streaked on LA and LA + TET 25 mg/L (to confirm loss of the plasmid).

#### 9.7 Identification and confirmation of mutants

##### 9.7.1 Identification and confirmation of deletion mutants

Deletion mutants should be TET sensitive and were identified by their PCR product size after performing colony PCR using DreamTaq DNA polymerase following the manufacturer's protocol (initial denaturation was 6 – 10 min). Primer pairs, annealing temperatures and expected product sizes are shown in below.

| Gene | Primer pair | $T_A$ (°C) | Size WT | Size mutant |
|---|---|---|---|---|
| *cpxS* | SHP 1105 + SHP 1120 | 59 | 2681 bp | 1358 bp |
| *cpxP* | SHP 1173 + SHP 1174 | 51 | 1951 bp | 1510 bp |
| *cpxR* | SHP 1165 + SHP 1166 | 55 | 2117 bp | 1439 bp |

Cryo-cultures of identified deletion mutants were prepared. For final confirmation of the mutation, the PCR products were purified with the *GeneJET PCR Purification Kit* or the *GeneJET Gel Extraction Kit* and Sanger sequenced by Eurofins Genomics (TubeSeq Service). The primers used for sequencing are listed below and samples were prepared according to company guidelines. Sequences were analysed with NCBI BLAST (24, 25).

| **Gene** | **Primers** |
|---|---|
| *cpxS* | SHP 1105, SHP 1120, SHP 1123, SHP 1124 |
| *cpxP** | SHP 1166, SHP 1105 |
| *cpxR** | SHP 1167, SHP 1168 |

* only primers inside the construct used for sequencing

#### 9.7.2 Identification and confirmation of *cpxS* SNP mutants

In contrast to deletion mutants, SNP mutants could only be identified by sequencing. Mutations in *cpxS* appear to come along with increased β-lactam resistance (23, 26, 27). Thus, CAR resistance might enable phenotypic differentiation between *cpxS* mutant and WT clones. Therefore, clones picked from TYS10 plates were not only re-streaked on LA and LA + TET 25 mg/L plates but also on LA + CAR (75 - 100 mg/L) plates to identify potential mutant clones. Cryo-cultures of clones of interest (both CAR resistant and susceptible) were prepared. For final identification of the desired *cpxS* mutants it was necessary to perform PCR with a proof-reading DNA polymerase to enable detection of SNPs. Colony PCR of candidate clones with Phusion DNA polymerase turned out to be challenging. Thus, genomic DNA was isolated from candidate clones and used as template DNA for PCR with Phusion DNA polymerase following the manufacturer's protocol (Buffer: Phusion GC Buffer, Primers: SHP 1105 & SHP 1120, $T_A$: 68.1 °C, exp. size: 2681 bp). The products were purified with the *GeneJET PCR Purification Kit* and Sanger sequenced by Eurofins Genomics (TubeSeq Service) or by the Institute of Clinical Molecular Biology in Kiel. The primers used for sequencing are listed below. Sequences were analysed with NCBI BLAST (24, 25) and the results confirmed the hypothesis of *cpxS* mutants being more resistant to CAR.

| **Desired mutant** | **Primers** |
|---|---|
| CpxS G188S | SHP 1139, SHP 1140 |
| CpxS DES81_83G | SHP 1141, SHP 1142 |

After final confirmation of the desired mutants (deletion and SNP mutants), all cryo-cultures of *E. coli* JM109 transformants and *P. aeruginosa* merodiploids were discarded. Only cryo-cultures of by sequencing confirmed mutants were kept.

### 10. *Detailed protocol for generation of inducible CpxS overexpression mutants*

Inducible CpxS overexpression mutants were generated using a restriction digest – ligation set up. The open reading frame of *cpxS* was cloned into the expression vector pME6032 containing an IPTG-inducible *LacI* - $P_{tac}$ expression system and a tetracycline resistance gene (28). CpxS overexpression mutants were generated in PA14 wild-type and *cpxS* deletion background. The detailed protocol for the generation of those strains is described below

#### 10.1 Preparation of the insert

First, primers that amplify the *cpxS* open reading frame (ORF) from PA14 and introduce restriction sites (upstream: *Eco*RI; downstream: *Xho*I) were designed. In addition, primers for the confirmation of the final construct were designed. A list of all primers used for the construction of CpxS overexpression mutants is shown below:

| **ID** | **Primer** | **Sequence 5' → 3'** |
|---|---|---|
| SHP 1131 | cpxS_F | actgtGAATTCatgcgttcactcttc |
| SHP 1132 | cpxS_R | actgtCTCGAGttattcggtcgcctt |
| SHP 1133 | pME6032-seq_F | cggctcgtataatgtgtgga |
| SHP 1134 | pME6032-seq_R | gatcgaaatccagatccttg |

The *cpxS* ORF was amplified from PA14 DNA with Phusion DNA polymerase following the manufacturer's protocol (Primers: SHP 1131 & SHP 1132, $T_A$: 66.6 °C). The size of the PCR product (~1350 bp) was checked by performing agarose gel electrophoresis. The PCR product was purified by pooling 2 PCR reactions followed by gel extraction (*GeneJET Gel Extraction Kit*).

#### 10.2 Preparation of the vector

The expression plasmid pME6032 was isolated from *E. coli* DH5a with the *QIAprep Spin Miniprep Kit*. The isolated plasmid was transformed into PA14 according to the protocol from Choi *et al.* (29). Electroporation was performed using the pre-set program *EC2* (1 pulse, 2.5 kV) of a *MicroPulser Electroporator* (Bio-Rad) and electroporation cuvettes with a gap width of 2 mm. Plasmid input concentration was 50 ng and recovery time was 1 h. Electroporated cells were plated on LA + TET 100 mg/L plates to select for transformants and incubated overnight at 37 °C. The next day, some transformants were pure-streaked on LA + TET 100 mg/L plates and colony PCR was performed to confirm the presence of the plasmid pME6032 using DreamTaq DNA polymerase following the manufacturer's guidelines (Primers: SHP 1133 and SHP 1134, initial denaturation: 6 min, $T_A$: 48.2 °C). The size of the PCR product (~160 bp) was checked by agarose gel electrophoresis and cryo-cultures of confirmed transformants (*P. aeruginosa* strain PA14: $OE_{empty}$) were prepared. For further steps pME6032 isolated from *P. aeruginosa* (*QIAprep Spin Miniprep Kit*) was used.

#### 10.3 Restriction digest and ligation

A restriction digest of purified pME6032 (vector) and purified *cpxS*-fragment (insert) with *Eco*RI (Thermo Scientific$^{TM}$; FD0274) and *Xho*I (Thermo Scientific$^{TM}$; FD2694) was performed. The vector was additionally digested with *Nco*I (Thermo Scientific$^{TM}$; FD0573) to cut the excised chunk and thus increase the chance that the chunk is lost during the purification step prior to ligation (the size cut-off is 25 bp, according to the manufacturer). Digest reactions were composed following the manufacturer's guidelines and incubated in a water bath at 37 °C for 3 h followed by 5 min heat inactivation at 80 °C. Digested DNA was purified with the *GeneJET PCR Purification Kit* (vector) or the *GeneJET Gel Extraction and DNA Cleanup Micro Kit* (Thermo Scientific$^{TM}$; K0831) (insert) according to the manual. Elution was performed with nuclease-free $H_2O$ instead of Elution Buffer. Afterwards, ligation reactions were set up using a molar vector to insert ratio of 1:3 (30 fmol : 90 fmol). The amount of DNA required was calculated with the NEBioCalculator. The ligation reactions were incubated at 16 °C in a thermocycler for 16 h and contained the following components:

| **Component** | **Amount** |
|---|---|
| Ligase Reaction Buffer (5 x) | 4 µL |
| digested vector | ~182 ng |
| digested insert | ~75 ng |
| T4 DNA Ligase (1 U/µL)* | 1 µL |
| nuclase-free $H_2O$ | to 20 µL |

* (Thermo Scientific$^{TM}$; 15224017)

Two ligation products were pooled and purified with the *GeneJET Gel Extraction and DNA Cleanup Micro Kit* according to the manufacturer's protocol. Instead of Elution Buffer nuclease-free $H_2O$ was used.

#### 10.4 Transformation in PA14 and confirmation of mutants

Purified ligation product was transformed into PA14 by electroporation (as described above). Input DNA amounts varied between approximately 150 and 540 ng. Transformants were pure-streaked on LA + TET 100 mg/L plates and colony PCR was performed to confirm the presence of the pME6032 plasmid with the insert as described above. The size of the PCR products was checked by performing agarose gel electrophoresis. Expected fragment sizes were ~1500 bp for the plasmid with the inserted *cpxS*-fragment (pME6032-CpxS) and ~160 bp for the empty plasmid. A subset of PCR products with the correct size was purified with the *GeneJET PCR Purification Kit* according to the manual and Sanger sequenced by Eurofins Genomics (TubeSeq Service) with the primers SHP 1133 and SHP 1134. Sequences were analysed with BLAST (24, 25). Finally, by sequencing confirmed transformants (*P. aeruginosa* strain PA14: CpxS-OE) were used to prepare cryo-cultures.

**10.5 Generation of a CpxS overexpression mutant in Δ*cpxS* background**

The plasmid pME6032-CpxS was isolated from *P. aeruginosa* PA14: CpxS-OE (*QIAprep Spin Miniprep Kit*) and transformed into Δ*cpxS* by electroporation as described above. Transformants were pure-streaked on LA + TET 100 mg/L plates. The next day, colony PCR was performed as described above to confirm presence of the plasmid. The size of PCR products (~1500 bp) was checked by agarose gel electrophoresis and cryo-cultures of clones containing the plasmid (*P. aeruginosa* strain Δ*cpxS*: CpxS-OE) were prepared.
